# Pervasive Antagonistic Interactions Among Hybrid Incompatibility Loci

**DOI:** 10.1101/090886

**Authors:** Rafael F. Guerrero, Christopher D. Muir, Sarah Josway, Leonie C. Moyle

## Abstract

Species barriers, expressed as hybrid inviability and sterility, are often due to epistatic interactions between divergent loci from two lineages. Theoretical models indicate that the strength, direction, and complexity of these genetic interactions can strongly affect the expression of interspecific reproductive isolation and the rates at which new species evolve. Nonetheless, empirical analyses have not quantified the frequency with which loci are involved in interactions affecting hybrid fitness, and whether these loci predominantly interact synergistically or antagonistically, or preferentially involve loci that have strong individual effects on hybrid fitness. We systematically examined the prevalence of interactions between pairs of short chromosomal regions from one species (*Solanum habrochaites*) co-introgressed into a heterospecific genetic background (*Solanum lycopersicum*). We used lines containing pairwise combinations of 15 chromosomal segments from *S. habrochaites* crossed into the background of *S. lycopersicum* (*i.e*., 95 double introgression lines). We compared the strength of hybrid incompatibility (either pollen sterility or seed sterility) expressed in each double introgression line to the expected additive effect of its two component single introgressions. We found that: epistasis was common among co-introgressed regions; epistastic effects for hybrid dysfunction were overwhelmingly antagonistic (i.e., double hybrids were less unfit than expected from additive single introgression effects); and, epistasis was substantially more prevalent in pollen fertility compared to seed fertility phenotypes. Together, these results indicate that higher-order interactions frequently contribute to postzygotic sterility barriers in these species. This pervasive epistasis leads to the decoupling of the patterns of accumulation of isolation loci and isolation phenotypes, and is expected to attenuate the rate of accumulation of hybrid infertility among lineages over time (*i.e*., giving diminishing returns as more reproductive isolation loci accumulate). This decoupling effect might also explain observed differences between pollen and seed fertility in their fit to theoretical predictions of the accumulation of isolation loci, including the ‘snowball’ effect.

**AUTHOR SUMMARY:** A characteristic feature of new species is their inability to produce fertile or viable hybrids with other lineages. This post-zygotic reproductive isolation is caused by dysfunctional interactions between genes that have newly evolved changes in the diverging lineages. Whether these interactions occur between pairs of divergent alleles, or involve more complex networks of genes, can have strong effects on how rapidly reproductive isolation—and therefore new species—evolve. The complexity of these interactions, however, is poorly understood in empirical systems. We examined the fertility of hybrids that carried one or two chromosomal regions from a close relative, finding that hybrids with two of these heterospecific regions were frequently less sterile than would be expected from the joint fitness of hybrids that have the same regions singly. This ‘less-than-additive’ effect on hybrid sterility was widespread (observed in 20% of pairwise combinations), and especially pronounced for male sterility. We infer that genes contributing to male sterility form a more tightly connected network than previously thought, implying that reproductive isolation is evolving by incremental dysfunction of complex interactions rather than by independent pairwise incompatibilities. We use simulations to illustrate these expected patterns of accumulation of reproductive isolation when it involves highly interconnected gene networks.

## INTRODUCTION

Intrinsic postzygotic isolation (hybrid inviability and sterility that occurs independently of the environment) is often due to deleterious genetic interactions between loci that have functionally diverged during the evolution of new species (*i.e*., Dobzhansky-Muller incompatibilities, or DMIs; [1]). Several models of this process assume that individual DMIs are due to epistasis between one locus in each diverging lineage (*i.e*., pairwise genetic interactions), and that each DMI contributes independently to the expression of hybrid incompatibility between diverging species (although some models can relax the first assumption; see [2-4]). If hybrid dysfunction is due to more complex interactions among loci, however, there can be important consequences for the temporal accumulation of species barriers and the number of loci required to complete speciation [4-6]. In the simplest case, if epistasis between different hybrid incompatibility loci is antagonistic (*i.e*., if the combined effect of two different DMIs is less than expected based on their individual effects on hybrid incompatibility), then a greater time to speciation is expected and correspondingly more loci are required. Conversely, if epistasis between different conspecific loci is typically synergistic (*i.e*., the combined effect on hybrid incompatibility is greater than expected based on individual effects), fewer DMIs will be required for the expression of complete reproductive isolation, with a correspondingly shorter time to speciation. In the latter case, although the initial observation of incompatibility phenotypes requires more than one substitution per lineage, hybrid incompatibility can be expressed rapidly between two diverging species once substitutions have begun to accumulate. The prevalence of epistasis, and whether this epistasis is synergistic or antagonistic, can therefore be critical in determining patterns and rates of evolution of isolation between diverging lineages by governing the accumulation dynamics of alleles that contribute to species barriers (*e.g*., [4, 5]; and see below).

Empirically, however, very little is known about the nature of epistasis among different loci contributing to hybrid incompatibility (‘complex epistasis’, c.f. [7]) and whether these interactions typically act to enhance or retard the expression of hybrid incompatibility between species. Some evidence suggests that non-additive interactions affecting the fitness of hybrids might be common. In *Drosophila*, for example, quantitative trait loci (QTL) for male sterility show evidence of complex epistasis. Several individual genomic regions are simultaneously required for the expression of some male sterility phenotypes (*e.g*., [8]; other evidence is reviewed in [1]), consistent with synergism (*i.e*., greater-than-additive effects) between conspecific loci for the expression of hybrid sterility. However, epistasis has been difficult to assess in many genetic analyses of quantitative trait loci because early recombinant generations have low power to detect these interactions [9]. The most promising method of directly estimating epistasis among loci is to subtract background effects by isolating two individual QTL in an otherwise isogenic background (*e.g*., [10]). For example, this approach revealed less-than-additive effects for quantitative and ecophysiological traits in tomato, consistent with antagonistic interactions between different conspecific loci [11, 12]. Similarly, studies in microorganisms have used serial pairwise combinations of target loci to examine the size and direction of pairwise epistasis on the phenotypic effects of, for example, double deletion strains in yeast [13], and metabolic flux mutants in yeast and *E. coli* [14]. Nonetheless, the serial combination of many pairs of conspecific loci has yet to be used to assess prevalence and direction of interactions influencing hybrid incompatibility phenotypes [15].

The goal of this study was to assess the prevalence and direction of genetic interactions between different chromosomal regions from one species when combined pairwise in the background of a second species (Figure S1), focusing specifically on the consequences of pairwise conspecific epistasis for the expression of hybrid incompatibility. We studied fifteen chromosomal regions (Table 1), drawing from a set of 71 introgression lines (ILs) between two plant species in the genus *Solanum* Section *Lycopersicon* (the tomato clade) [16]. Each IL contains a unique short chromosomal region from the wild species *S. habrochaites* (‘*hab*’) introgressed into the otherwise isogenic genetic background of the domesticated tomato, *S. lycopersicum* (‘*lyc*’) ([16]; see [17] for a previous summary). The fifteen ILs included in our study, several of which have effects on either pollen or seed sterility ([17], and see Results), were previously chosen and crossed to generate nearly 100 double introgression lines (DILs; [18]). Here, we draw on this collection of DILs to study the effect of interactions among conspecific introgressed regions on the expression of known hybrid incompatibility phenotypes. Specifically, we compared fertility phenotypes in homozygous DILs to those of their corresponding parental ILs to infer frequency, magnitude and direction of epistasis among conspecific introgressions.

**Table 1:**
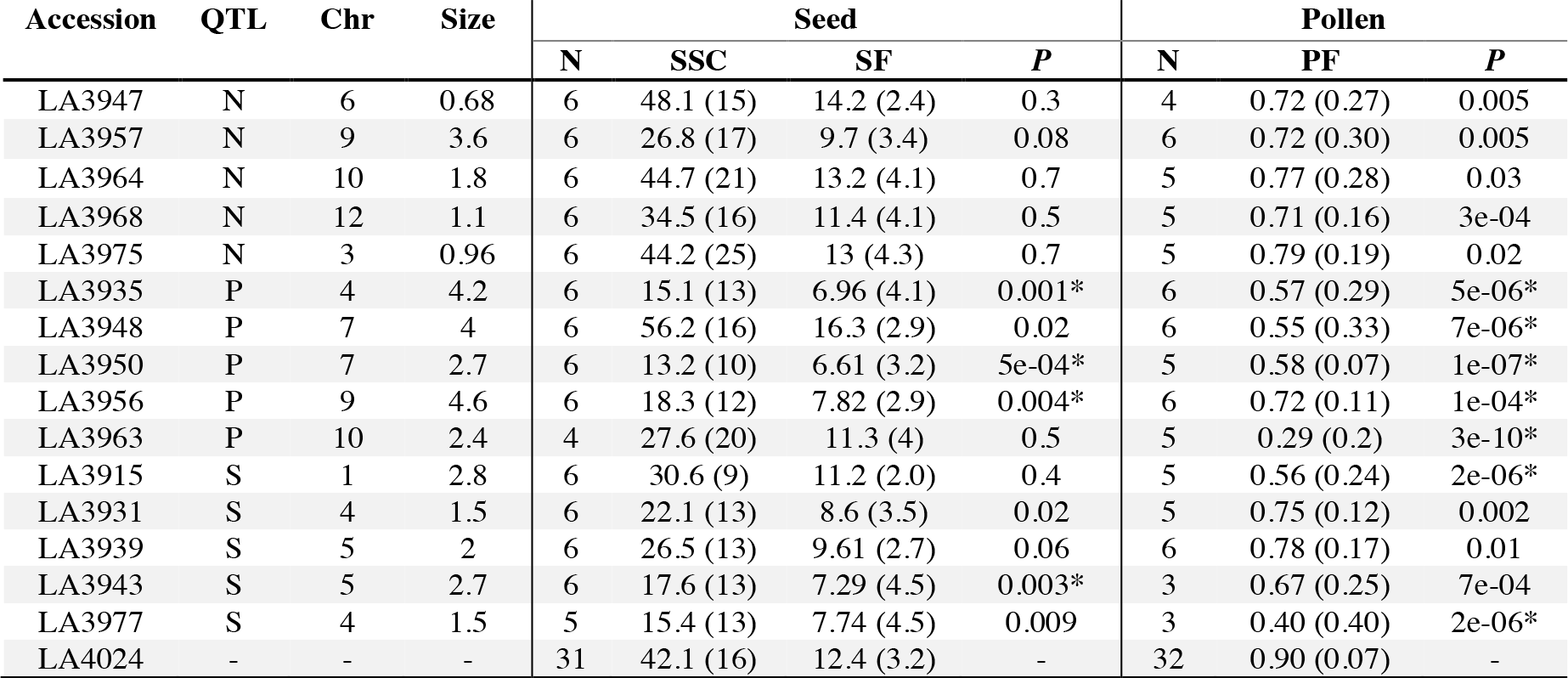
Introgression lines (ILs) phenotyped and used as parents to generate double-introgression lines. Accession identifiers from the Tomato Genome Resource Center (tgrc.ucdavis.edu). QTL: Prior evidence of IL carrying QTL for pollen sterility (P), seed sterility (S) or none (N). Chr: Chromosomal location of the introgression (out of 12 chromosomes in the *lyc* genome). Size: length of introgressed region, as a percentage of the *lyc* genome. Seed: Sample size (N), mean (and standard deviation) for self-seed count (SSC) and pollen-corrected seed fertility (SF). Pollen: Sample size (N), mean (and standard deviation) for proportion of fertile pollen (PF). *P*-values are from normal and quasi-likelihood binomial general linear models (for seed and pollen, respectively). (*) The IL has a significant individual effect on the corresponding phenotype.

| Accession | QTL | Chr | Size | Seed |  |  |  | Pollen |  |  |
| --- | --- | --- | --- | --- | --- | --- | --- | --- | --- | --- |
|  |  |  |  | N | SSC | SF | *P* | N | PF | *P* |
| LA3947 | N | 6 | 0.68 | 6 | 48.1 (15) | 14.2 (2.4) | 0.3 | 4 | 0.72 (0.27) | 0.005 |
| LA3957 | N | 9 | 3.6 | 6 | 26.8 (17) | 9.7 (3.4) | 0.08 | 6 | 0.72 (0.30) | 0.005 |
| LA3964 | N | 10 | 1.8 | 6 | 44.7 (21) | 13.2 (4.1) | 0.7 | 5 | 0.77 (0.28) | 0.03 |
| LA3968 | N | 12 | 1.1 | 6 | 34.5 (16) | 11.4 (4.1) | 0.5 | 5 | 0.71 (0.16) | 3e-04 |
| LA3975 | N | 3 | 0.96 | 6 | 44.2 (25) | 13 (4.3) | 0.7 | 5 | 0.79 (0.19) | 0.02 |
| LA3935 | P | 4 | 4.2 | 6 | 15.1 (13) | 6.96 (4.1) | 0.001* | 6 | 0.57 (0.29) | 5e-06* |
| LA3948 | P | 7 | 4 | 6 | 56.2 (16) | 16.3 (2.9) | 0.02 | 6 | 0.55 (0.33) | 7e-06* |
| LA3950 | P | 7 | 2.7 | 6 | 13.2 (10) | 6.61 (3.2) | 5e-04* | 5 | 0.58 (0.07) | 1e-07* |
| LA3956 | P | 9 | 4.6 | 6 | 18.3 (12) | 7.82 (2.9) | 0.004* | 6 | 0.72 (0.11) | 1e-04* |
| LA3963 | P | 10 | 2.4 | 4 | 27.6 (20) | 11.3 (4) | 0.5 | 5 | 0.29 (0.2) | 3e-10* |
| LA3915 | S | 1 | 2.8 | 6 | 30.6 (9) | 11.2 (2.0) | 0.4 | 5 | 0.56 (0.24) | 2e-06* |
| LA3931 | S | 4 | 1.5 | 6 | 22.1 (13) | 8.6 (3.5) | 0.02 | 5 | 0.75 (0.12) | 0.002 |
| LA3939 | S | 5 | 2 | 6 | 26.5 (13) | 9.61 (2.7) | 0.06 | 6 | 0.78 (0.17) | 0.01 |
| LA3943 | S | 5 | 2.7 | 6 | 17.6 (13) | 7.29 (4.5) | 0.003* | 3 | 0.67 (0.25) | 7e-04 |
| LA3977 | S | 4 | 1.5 | 5 | 15.4 (13) | 7.74 (4.5) | 0.009 | 3 | 0.40 (0.40) | 2e-06* |
| LA4024 | - | - | - | 31 | 42.1 (16) | 12.4 (3.2) | - | 32 | 0.90 (0.07) | - |

We found that complex interactions were common in both pollen and seed fertility phenotypes, although significantly more frequent for introgressions affecting pollen sterility. These observed interactions were frequently antagonistic, whereby the combined effect of pairwise introgressions produced a less severe effect on fitness than predicted from individual effects. For pollen sterility phenotypes, we found that some chromosomal regions were considerably more prone to interaction than others: most interactions occurred among introgressions with significant individual effects. In contrast, we found no evidence of a similar pattern for seed sterility.

This pervasive antagonism has critical implications for the predicted rate and pattern of accumulation of hybrid incompatibility between these lineages, and for future empirical studies of the evolution of post-zygotic isolation. We found unexpectedly abundant interactions among genes underlying the phenotypes studied, which implies a violation of key assumptions—regarding the independence among different pairs of incompatibility loci—made by many speciation models. We propose that this high connectivity among loci that contribute to pollen sterility could explain the observed flattened accumulation of (pairwise) pollen incompatibilities in this clade [18]. In addition to explaining this apparent lack of “snowball” effect, our results also imply that standard QTL mapping approaches can underestimate the number of incompatibility-causing interactions present in systems with high levels of epistasis.

## METHODS

We assessed fertility phenotypes for 110 unique genotypes: the isogenic parental *lyc*, 15 homozygous introgression lines (ILs), and 95 double-introgression lines (DILs), each of which were generated previously. Monforte et al. [16] constructed a library of over 70 introgression lines that cover 85% of the *hab* genome in the background of *lyc*. Each line contains a single, marker delimited chromosomal region from *hab* introgressed into the background of *lyc*. Using this IL library, we had detected 8 and 5 QTL for hybrid pollen and seed sterility, respectively, between these species [17]. Based on these results, Moyle & Nakazato [15] chose 15 ILs (covering about 36% of the donor genome) to use in the construction of DILs. Ten of these lines carried QTL for hybrid incompatibility (five with significant pollen, and five with significant seed, sterility effects) when compared to pure parental genotypes. The remaining five lines contain *hab* introgressions with no previously detected phenotypic effect on pollen or seed sterility; these additional lines were chosen based on their distribution in the genome (each was located on a different chromosome), and to be roughly comparable in terms of introgression length (as a percent of *hab* genome introgressed) as the partially sterile lines (Table 1). Nonetheless, because the detection of sterility effects can be sensitive to environmental conditions and dependent on experimental power, all 15 lines were reassessed for sterility effects in the current experiment (see Results).

To generate the DILs, Moyle & Nakazato [15] performed a full diallel cross among the 15 ILs to combine each introgression with each other introgression (producing a total of 105 unique pairwise combinations; Figure S1). Heterozygote F_1_ DILs from each of these 105 pairwise IL combinations were selfed to generate F_2_ seeds, and up to 120 F_2_s in each DIL family were grown and genotyped for two markers, one located in each introgressed region. From these families, individuals that were confirmed to carry two *hab* alleles at both introgression locations (i.e., double *hab* homozygotes) were retained. For the current experiment, these lines were selfed to produce double *hab* homozygous seed. Out of the 105 DILs, we excluded one with overlapping introgressions and one due to ambiguous genotyping. From the remaining 103 lines, seven failed to produce seed from self-fertilizations, despite repeated hand-pollinations; these genotypes were necessarily excluded from the experiment (see summary of missing data in Table S1). Note that we focus our analysis here solely on comparisons of fertility between ILs and DILs that are homozygous for *hab* introgressions. This is because the offspring of heterozygous NILs and DILs are expected to segregate for introgressions, preventing an equivalently direct comparison of fertility between these classes of lines. These genotypes and phenotypes are therefore not analyzed here.

### Assessment of fertility phenotypes

Each unique genotype was grown and assessed in replicate (total experiment size = 702 individuals; Table S2) in a fully randomized common garden (greenhouse) experiment at the IU Biology greenhouse facility. Cultivation methods have been previously described [12]. Briefly, all experimental seeds were germinated under artificial lights on moist filter paper, transferred to soil post-germination (at the cotyledon stage), and repotted into 1gallon pots at 4-6 weeks post-germination. A total of 652 individuals reached reproductive maturity. Due to mortality and/or fecundity variation among individuals, we were able to collect pollen and seed fertility data from 569 and 620 individuals, respectively (Table S2). Because of these inviability or fertility effects, not all DIL genotypes had biological replicates and so were not included in analyses. In total, we analyzed pollen and seed fertility data for 95 and 93 DILs, respectively (see Table S3, Results and Discussion).

Seed fertility (seeds per fruit) was determined by quantifying seed production from self-pollinated fruits (‘self-seed set’), as with previous QTL analyses [19]. At least two flowers per plant were allowed to produce fruit via selfing; when fruits did not develop automatically, flowers were self-pollinated manually to ensure that floral morphology was not responsible for preventing self-fertilization. Upon maturation, individual fruits were harvested, seeds extracted by hand, and seed fertility determined by counting the number of visible seeds from each fruit. Average seed per fruit for each plant was used to generate self-seed set estimates for each introgression genotype and the *lyc* control parent.

Pollen fertility (quantified as the proportion of fertile pollen) was estimated on two unopened flowers on each plant, as previously described [17]. Briefly, all pollen (the entire anther cone) from each target flower was collected into lactophenol-aniline blue histochemical stain, homogenized, and a known sub-sample of homogenate used to count inviable and viable pollen grains using microscopy. Pollen inviability was indicated by the absence of a stained cytoplasm, a conservative measure of pollen infertility [20].

Seed set phenotypes may be affected by three components: ovule viability, pollen fertility, and gamete compatibility in the zygote. Therefore, pollen sterility in our lines may influence estimates of seed sterility. In fact, we found a weak but significant correlation between pollen fertility and self-seed set across our experimental lines (*P*=0.0014; negative-binomial GLM). However, pollen fertility explained only about 5% of variation in seed fertility (pseudo-R^2^ = 0.046), which is consistent with previous studies [17, 19]. Nonetheless, we removed this small pollen effect by carrying out our analyses of self-seed set on their residuals from the regression on pollen fertility (including the genotype as a random effect in a GLM). For completeness, we carried out the same analyses on the uncorrected self-seed set values; our results do not differ qualitatively between these analyses (Figure S2) but, as the ‘pollen-corrected’ data is more conservative with respect to inferring sterility effects, we report these results here.

### Analysis

#### Estimation of sterility in ILs

To quantify the individual effect of introgressions on hybrid fitness we tested for differences between the phenotypes of the ILs and those of the background parent, *lyc*, using generalized linear models (GLMs) with a single fixed effect. Pollen fertility was tested assuming a quasi-binomial error distribution appropriate for proportion data. Seed fertility data were analyzed assuming a normal error distribution (after pollen-correction, see above). Fitness of each IL was tested independently against *lyc*, allowing slightly different significance thresholds for each fitness component. For corrected self-seed set data, we took ‘sterile’ ILs at a false discovery rate (FDR) of 5% (which implied *P* < 0.005; Table 1). We were slightly more stringent with significance for pollen fertility, for which we took ‘sterile’ ILs at a FDR of 1% (which corresponded to *P* < 10^−4^). The significance threshold of individual IL effects on sterility does not affect our key findings (see Results, Figure S3).

#### Inference of epistatic effects in DILs

We used a likelihood maximization framework to infer the presence and strength of epistasis from DIL phenotypic data. In our likelihood approach fitness can be either additive (where fitness effects in DILs are the sum of the fitness effects of the two parental ILs) or multiplicative (in which DIL fitness is the product of parental IL fitness). These common alternative fitness models were included to allow us to evaluate epistatic interactions for each DIL (*i.e*., to compare non-epistatic ‐‐or null‐‐ and epistatic models, as outlined below) without having to assume *a priori* precisely how introgressions combine to affect fitness in each DIL genotype. Given this, we did not test support for choice between additive and multiplicative models of fitness. (Indeed, there was little evidence for significant differences in the fit of these models; see Results.) Instead, we estimated statistical support for the choice between null vs. epistatic models, in line with our primary goal to assess evidence for epistastic interactions.

First, we define fitness in DILs and their IL parents as the phenotype (pollen or seed fertility) relative to the mean phenotype in *lyc*. For each DIL carrying introgressions *i* and *j*, we assume that relative fitness, *w*_*ij*_, is normally distributed with mean and variance, *μ*_(*ψ*)_ and 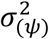, determined by the fitness model *ψ* (={*a, p*}), relative fitness of the parental ILs (*w*_*i*_ and *w*_*j*_) and ε_*ij*_, the epistasis parameter. Under the additive fitness model, *μ*_(*a*)_ and 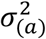, are the mean and variance of the vector 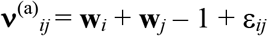, where **w**_*i*_ and **w**_*j*_ are vectors holding *k* observed values for *w*_*i*_ and *w*_*j*_ (that is, there are *k* biological replicates of each parental IL). Similarly, under the multiplicative fitness model the parameters *μ*_(*p*)_ and 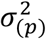 are the mean and variance of the vector 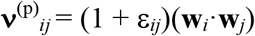, with the underlying simplification that the product of two normally distributed random variables (*w*_*i*_ and *w*_*j*_) is approximately normal for parameters *μ*_(*p*)_ = *O*(1) and 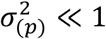.

The likelihood function for a single DIL under model *ψ* is 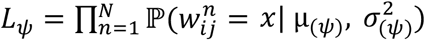, the product of the normal probability density over *N* observations of relative fitness. In both models, the likelihood function has the epistasis term, ε_*ij*_, as a single free parameter.

We maximized likelihood for each model numerically (using the *mle* function from the stats4 package in R; [21]). To improve normality of our observations, we used arcsine transformed values for pollen fertility. Because we have different sample sizes for the phenotypes of parental ILs (*i.e*., vectors **w**_*i*_ and **w**_*j*_ have different lengths), we subsampled without replacement each set of observations to the minimum parental pair of each DIL (that is, took *k_m_* = min(*k*_*i*_, *k*_*j*_) elements from each vector). We repeated this subsampling 100 times, and took the average of *μ*_(*ψ*)_ and 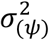 from all of them to obtain estimates that use all our data and minimize the effect of pairing observations from parental ILs.

To choose the best model, we compared the resulting maxima using the Bayesian Information Criterion (BIC; [22]). Additionally, to confidently infer cases of epistasis, we required that our best-fit epistasis model have a better BIC score than that of two null models (both additive and multiplicative fitness models with ε*_ij_* = 0). Finally, we obtained statistical support for our inference of epistasis through a likelihood-ratio test of the best-fit epistasis model against the best null model (assuming a chi-squared distribution of the log-likelihood surface and 1 degree of freedom). We applied a correction for multiple testing within each phenotype, taking significant epistatic interactions at a 1% FDR for both pollen and seed sterility. We visualized significant interactions between ILs as a network using the *igraph* [23] and *ggraph* packages [24]. In these networks, each node represents an IL (specifically, the introgressed chromosomal region within that IL), while edges represent an epistatic interaction between two ILs. All scripts used, written in R [21], are available in the Supplement (File S1).

Our approach is similar to the one described by Gao *et al*. [25] for the classification of epistasis. In contrast to that previous work, however, we have not considered a third fitness model (the ‘minimum’ model) in our reported estimations. The ‘minimum’ model aims to assess statistical independence among mutations, however it leads to difficulty detecting certain types of biological interactions (*e.g*., ‘masking’ and ‘co-equal’ interactions; [26]), and might not be adequate when testing epistasis among loci with known similar functions [25]. Moreover, when our data are re-analyzed including this model, there is no change in the direction and only a slight decline in the frequency of significant epistatic interactions (Table S3).

For simplicity, throughout this manuscript we refer to individual effects of QTL and interaction effects between loci –although strictly discussing the phenotypic consequences of entire introgressions. The estimated length of some of our introgressions (Table 1) suggests that they could carry up to several hundred genes each. Indeed, we found a weak correlation between the size of an introgression and its total number of interactions (negative binomial GLM, *P* = 0.02, pseudo-R^2^=0.07; Figure S4).

Accordingly, our study more accurately is an analysis of the frequency and strength of *trans* interaction effects between short conspecific chromosomal regions and is unable to differentiate, for example, the single versus combined phenotypic effects of multiple genes within any given introgression. Depending upon genetic structure of these QTL at finer chromosomal scales, the specific genetic interpretation of types and dynamics of substitutions would change. We note, however, that most observations of complex epistasis outside of microbial studies have come from examining *trans* interactions between chromosomal regions rather than individual genes (*Drosophila*, [27]; *Tigriopus*, [28]; *Mimulus*, [29]). In addition, prior QTL mapping studies in this and other *Solanum* species pairs indicate that the genetic basis of hybrid incompatibility is modestly, not highly, polygenic [17, 19]; therefore, unless individual loci underlying sterility QTL are unusually (and unexpectedly) highly clustered within the genome, these data suggest most introgressions do not harbor multiple different sterility-causing loci.

## RESULTS

We found three prevalent patterns of conspecific epistasis for hybrid incompatibility. First, epistatic interactions between conspecific loci are common, especially among introgressions with larger individual effects on fertility (Figures 1 and 2). Second, these interactions are predominantly antagonistic (*i.e*., less-than-additive) in their effects on hybrid incompatibility phenotypes. Third, both these patterns are significantly more pronounced for pollen than seed sterility effects.

**Figure 1:**
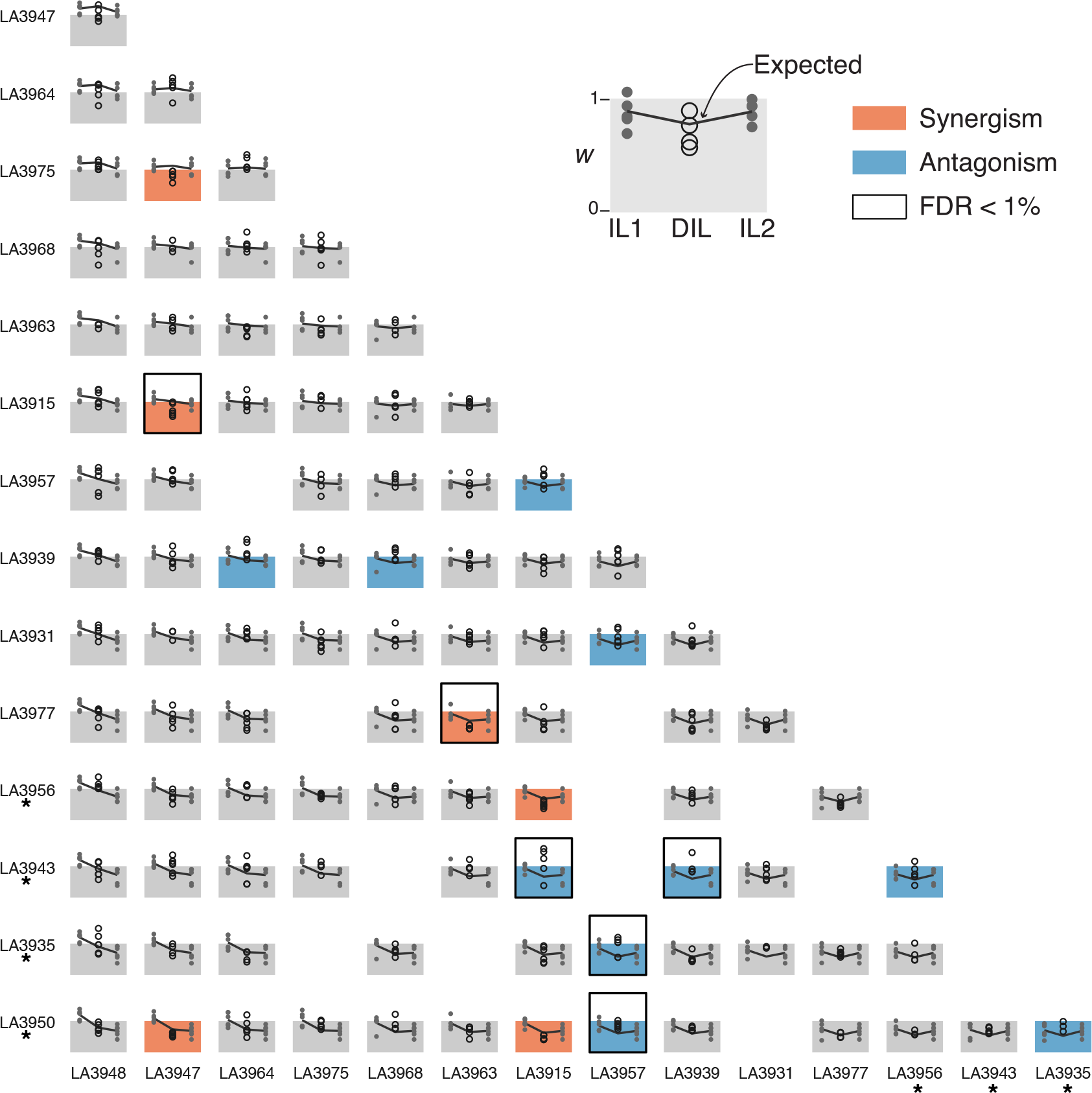
Self-seed set, relative to *S. lycopersicum*, for all parental introgression lines (ILs) and double-introgression lines (DILs). Parental ILs (across rows and columns) are arranged by their individual effect on the phenotype (larger effects down and to the right). Parental ILs with significant individual fitness effects are marked with an asterisk. Each panel shows the observed values for two parents (small circles) and their corresponding DIL (large circles, center in each panel). Lines connect the observed mean phenotypes for each parental IL and the expected mean phenotype for the DIL. The background of each panel spans from 0 to 1 along the vertical axis (1 corresponds to the mean fitness value of *S. lycopersicum*). Accented panels show significant support (*P* < 0.01) for epistasis (antagonistic interactions = blue panels; synergistic = red panels). Black bordered panels indicate highly significant interactions (FDR 1%).

**Figure 2:**
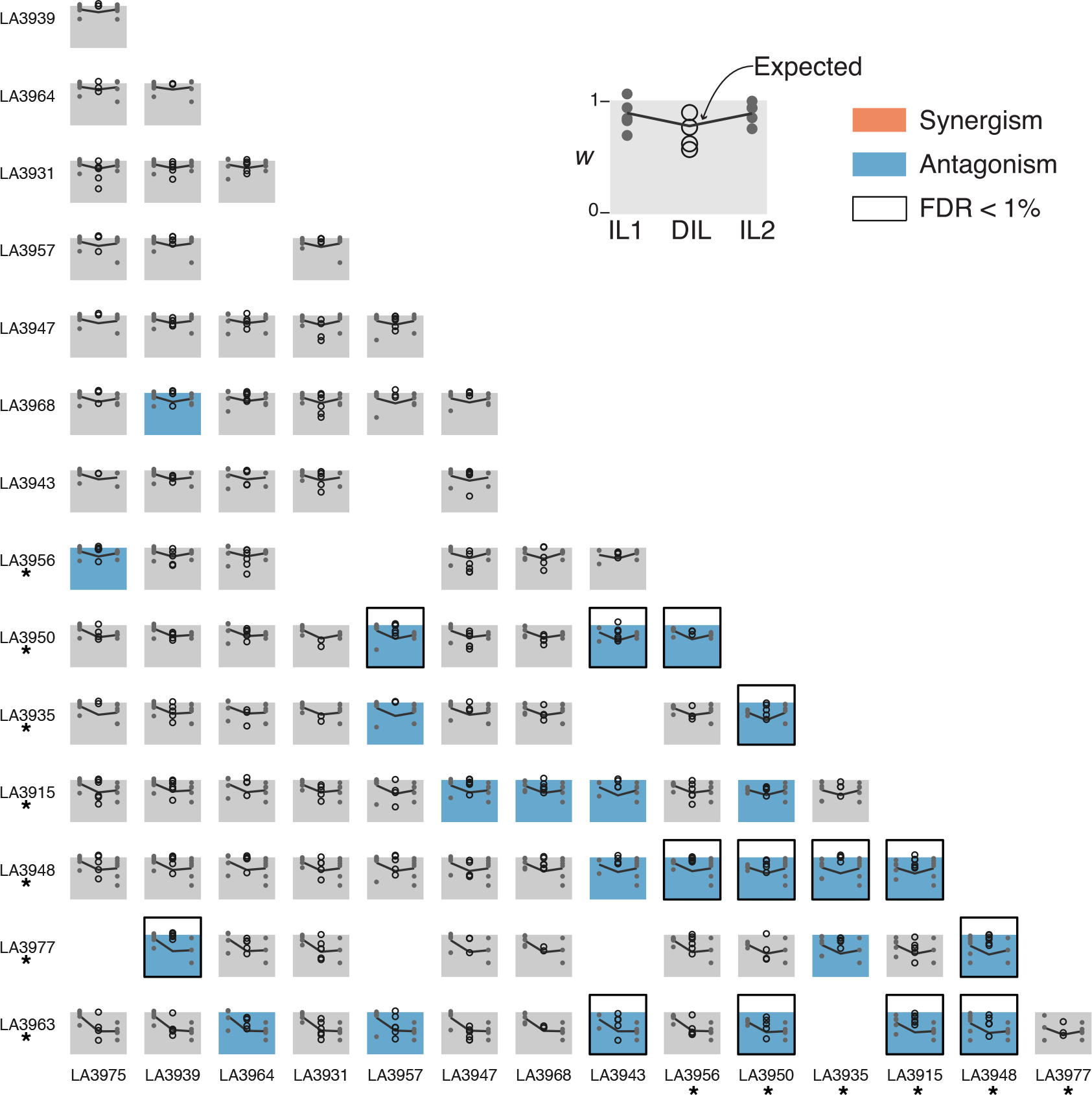
Proportion of fertile pollen, relative to *S. lycopersicum*, for all parental introgression lines (ILs) and double-introgression lines (DILs). All conventions as in Figure 2.

For seed sterility, we found evidence of fertility-affecting interactions between introgressions in about one fifth of DILs analyzed (Figure 1). Out of 93 DILs, 16 showed departures from fitness models of independence (*P* < 0.01, FDR=5.6%). Interactions were qualitatively more common for pollen fertility phenotypes (Figure 2), where 25 out of 95 analyzed DILs were best fit by epistatic fitness models (*P* < 0.01, FDR=3.4%). Remarkably, most of these interactions were antagonistic: in seed phenotypes, 10 DIL hybrids were less sterile than expected by their IL parents’ individual effects. This pattern was considerably more dramatic for pollen fertility, where all 25 detected epistatic interactions were antagonistic (Figure 2, Table S3).

We found little difference between pollen and seed sterility phenotypes in the magnitude of epistasis or the best fit fitness model (Table S3). Epistatic interactions caused, on average, a 37% excess in pollen fitness when compared to the fitness predicted from the joint effect of parental ILs. In seed sterility, we found similar effect sizes in synergistic and antagonistic interactions (33% and 45%, respectively). In most DILs with significant epistatic effects, an additive fitness model had a better fit than a multiplicative one (10 out of 16 in seed; 21 out of 25 in pollen); this was also true in DILs with no significant epistatic effects for seed (39 out of 77 DILs) but not for pollen phenotypes (16 out of 70). Note, however, that most differences in likelihood values between the two fitness models are probably not significant: only 6 cases (out of 188 analyzed DIL phenotypes) showed likelihood differences that would be significant in a chi-squared test (*i.e*., greater than 3 log-likelihood units, roughly *P* =0.01). This apparent bias in favor of an additive fitness model could be because this model predicts slightly larger variance than its multiplicative counterpart, resulting in a better fit even when there are no biologically meaningful differences in predicted means.

Epistatic interactions for fitness were common across the genome: most parental ILs were involved in at least one interaction with another conspecific introgression (all except *LA3948* for seed and *LA3931* for pollen fertility; Figures 1 and 2). In pollen phenotypes, however, interactions more frequently involved introgressions that had individual sterility effects. This bias becomes more apparent in highly significant interactions (14 DILs at FDR=1%; bordered panels in Figure 2), which always involved DILs with at least one pollen-sterile IL parent genotype; moreover, 10 of these 14 highly significant interactions involved two sterile ILs.

Individually, each of the observed interactions could be used to infer whether the incompatibility they participate in is pairwise or higher-order (Figure 3). A pairwise DMI could be inferred when an antagonistic interaction is observed between a sterile and a non-sterile IL (found in 4 cases for seed, and 4 for pollen phenotypes), since phenotype rescue in the DIL is consistent with complementation of loci carried by parental ILs. On the other hand, higher-order DMIs could be inferred from two types of interactions. First, all synergistic interactions (2 cases in seed phenotypes) imply introgressions that are part of a third-order (or higher) DMI in which a specific combination of alleles is necessary for sterility. Second, antagonistic epistasis between sterile ILs (the most common in our data; 10 cases, all in pollen phenotypes) implies an incompatibility involving at least three loci. We can rule out a pairwise DMI, which would not be consistent with the expectation that the complementing allele should be compatible with the heterospecific background (that is, there should be asymmetry of DMI allelic effects [30]). However, we cannot determine how these three or more loci interact: the parental ILs could carry loci involved in a single higher-order DMI, or the interaction could be the result of complex epistasis between two pairwise incompatibilities.

**Figure 3.**
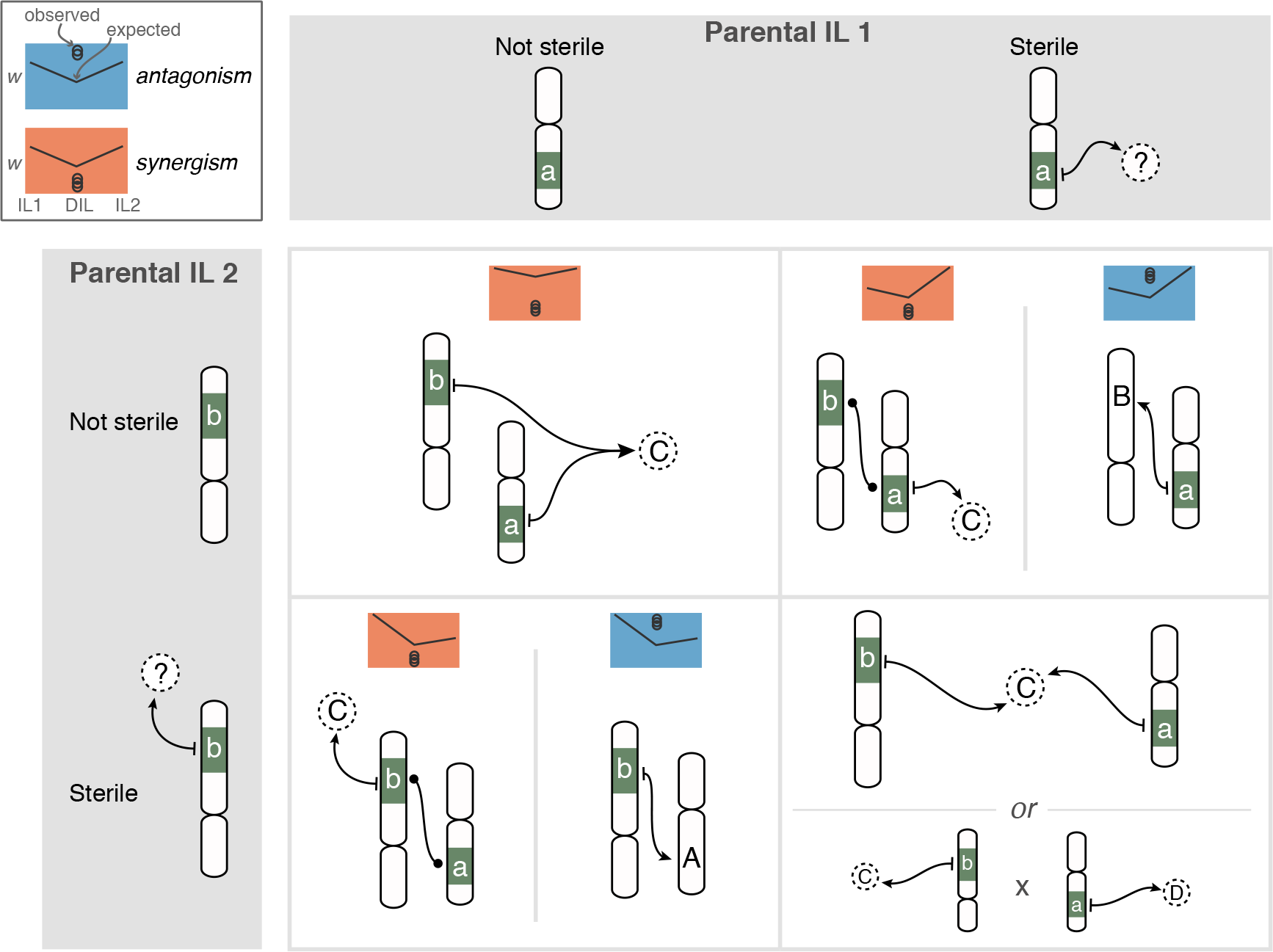
Possible DMI architecture inferred from observed epistasis in DILs. Synergistic epistasis (orange boxes), regardless of fitness of the parents, implies a higher-order DMI that involves *hab* alleles in both introgressions (a, b; green background) and at least one *lyc* locus at an unknown genomic location (C, dashed circles). Antagonistic interactions (blue boxes) between sterile and non-sterile ILs indicate a pairwise DMI (either a,B in the upper right or A,b in the lower left). Interactions, in any direction, between two sterile ILs suggests a higher-order DMI or complex epistasis between two DMIs (lower right panels).

Regardless of these individual inferences, the overall pattern of interactions we detect indicates there is a complex landscape of interdependence among sterility loci. This pattern can be visualized by representing the distribution of interactions as a network (Figure 4). In pollen, ILs formed a single interconnected cluster in which each line had, on average, 1.9 highly significant interactions. Sterile ILs for pollen had more interactions (*i.e*., a higher node degree; 3.5 on average) than non-sterile introgressions (0.5 on average). In contrast, the overall network of interactions in seed phenotypes was sparser than in pollen phenotypes, with three separate clusters of up to 4 ILs. The seed sterility network showed no direct connections between two sterile ILs, and sterile ILs had similar number of interactions than non-sterile ILs (0.8 on average for both groups). The number of interactions in which an IL was involved is not correlated with its mean effect size for seed sterility (unlike for pollen sterility; Figure S3), suggesting that the lower overall connectivity and reduced enrichment for individually sterile ILs we observed in seed phenotypes is robust to our arbitrary cut-off for significance of individual IL sterility (FDR of 1% for pollen and 5% for seed).

**Figure 4:**
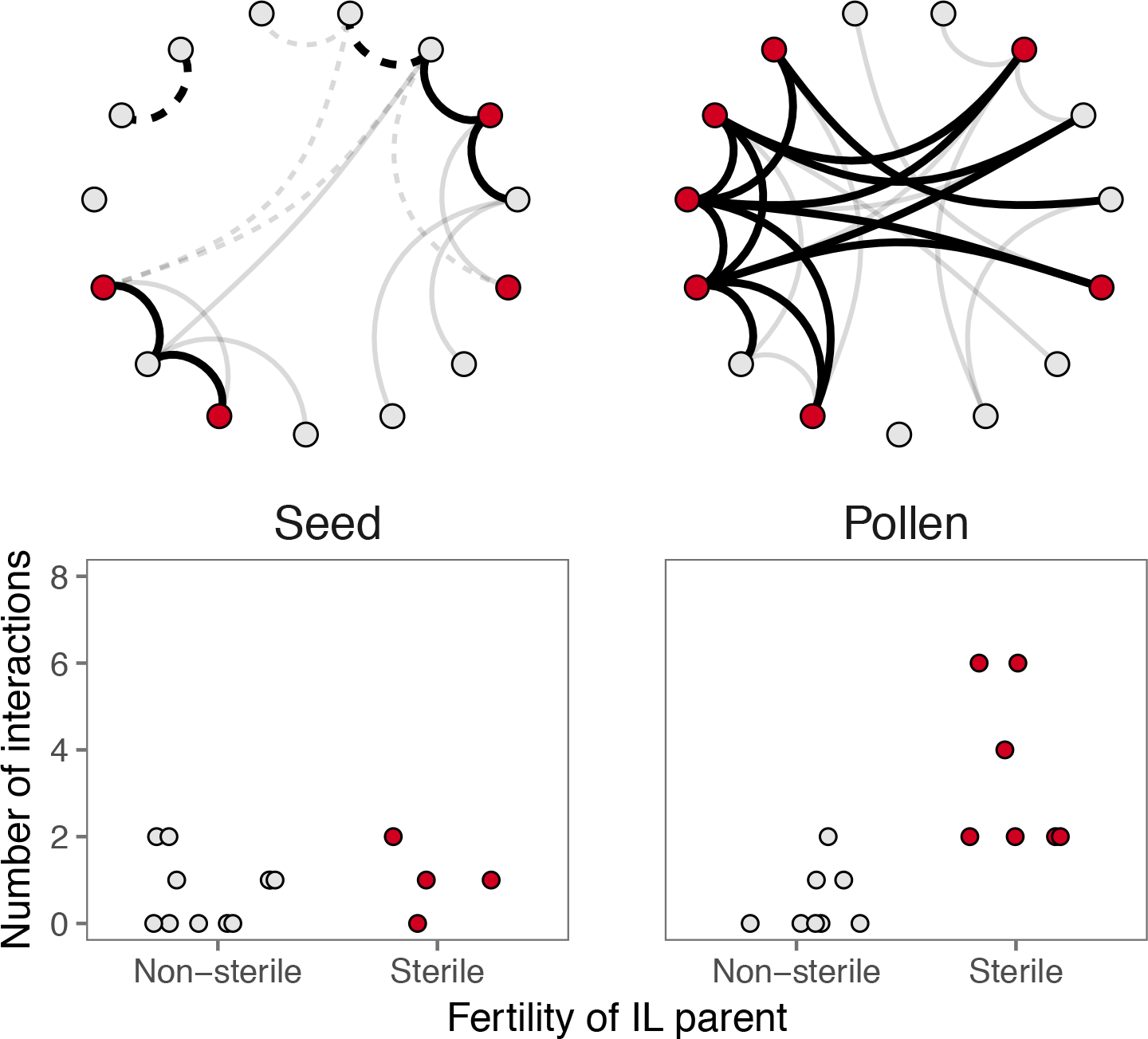
Above, the network representation of interactions in self-seed set (left) and proportion of fertile pollen (right). Parental ILs (nodes; red = sterile IL) are connected if they showed a highly significant interaction (FDR=1%). Solid lines denote antagonistic interactions; dashed lines mean synergistic interactions. Below, the relationship between parental IL fertility on the number of interactions (node degree) showed.

## DISCUSSION

By examining multiple pairwise combinations of conspecific loci, co-introgressed into a heterospecific background, we have shown evidence for pervasive non-independence among introgressions. This non-independence is likely due to a combination of pairwise interactions, higher-order incompatibilities, and complex epistatic effects, and suggests that hybrid phenotypes might frequently be the product of epistatic interactions among several factors. For pollen sterility phenotypes, we found that some chromosomal regions were more prone to interaction than others: most interactions occurred among introgressions with significant individual effects. In contrast, we found no evidence of a similar pattern for seed sterility. Finally, the observed interactions in pollen phenotypes were overwhelmingly antagonistic, whereby the combined effect of pairwise introgressions produced a less severe effect on fitness than predicted from individual effects. This observed pervasive antagonism could have important implications for understanding both the predicted patterns and the mechanism of accumulation of hybrid incompatibility between lineages, as well as for the detection of such loci in QTL mapping efforts.

Interactions were common in both pollen and seed fertility phenotypes, although considerably more frequent for introgressions affecting pollen sterility. Interactions in seed suggest at least two higher-order incompatibilities, composed of three and four loci (if we focus on highly significant edges in Figure 4). In the pollen network, ILs form a single cluster of highly significant interactions, and most of these interacting ILs are epistatic with at least two other introgressions (8 out of 10 ILs; Figure 4). Speculating on the genetic details underlying these patterns of interdependence is not straightforward. One extreme interpretation is that this large module represents a single incompatibility that involves ten loci (*i.e*., a tenth-order DMI). Alternatively, each of these antagonistic interactions could represent a pairwise hybrid incompatibility, which implies that pollen-sterility loci are frequently involved in more than one DMI (since many ILs show more than one interaction). This latter interpretation is also consistent with our observation that pollen sterile ILs show more interactions than non-sterile ones, since each interaction in which they engage could contribute to their observed fitness (*i.e*., ILs involved in more DMIs would, on average, show larger fitness effects). While our approach does not allow us to distinguish between these alternatives, our observations likely represent a mix of higher-order DMIs and loci involved in multiple pairwise incompatibilities. However, the involvement of loci in more than one hybrid incompatibility violates a central assumption in classical models of their evolutionary accumulation (an issue that we examine below).

The differences in connectivity between seed and pollen phenotypes could explain previously observed differences in the pattern of accumulation of hybrid incompatibilities (or ‘snowball’ effect) in this group. Moyle & Nakazato [18] showed that, while the accumulation of seed sterility QTL among increasingly divergent *Solanum* lineages follows the predicted snowball (*i.e*., it is faster than linear), this does not seem to be the case for QTL involved in pollen fertility. At first, this linear accumulation of pollen fertility QTLs may seem at odds with our observation of high connectivity for the phenotype, because theory predicts a faster increase in the number of incompatibilities with increasing connectivity among loci. In the simplest version of the snowball model, the number of incompatibilities is roughly proportional to the square of *K*—the number of genetic differences between two species, and *p*—the probability that any pair of these is incompatible [30]. Therefore, factors that increase *K* or *p* should increase the speed of the snowball. For instance, *K* could be elevated for phenotypes experiencing selective pressures, such as antagonistic coevolution, that result in sustained elevated rates of substitution [31, 32]. Similarly, the probability that any pair of genetic differences is incompatible (*p*) is elevated when there are more opportunities for incompatibilities to arise, such as when the phenotype is controlled by a highly connected network of genes. Given these predictions, our results would suggest that the number of detected pollen sterility loci should snowball faster than seed sterility loci (yet, they do not).

A possible explanation for this discordance lies in the assumption that the number of loci involved in incompatibilities represents an adequate proxy for the total number of incompatibilities. Because of the challenges of identifying underlying molecular loci, empirical studies have used the number of sterility QTL as an indicator of the number of DMIs, with the underlying assumption that each QTL is involved in only one (pairwise) incompatibility. Under a pairwise snowball model, each locus is involved in (*K*-1)*p* incompatibilities on average [3]. Since *p* (the probability of an incompatible pair) is assumed to be very small compared to *K*, such that (*K*-1)*p* << 1, each QTL is therefore likely to be involved in just one incompatibility. In cases with increased *p*, however, sufficient genetic differences could accumulate that (*K-*1)*p* ≥ 1 (*i.e*., loci are, on average, involved in one or more incompatibilities). After this “saturation” point, each new genetic difference will continue to generate (*K-*1)*p* new incompatible interactions, but the number of loci involved in hybrid incompatibility increases just by one. Note that many loci could be involved in more than one incompatibility before the system reaches saturation so that the deceleration of the snowball need not be abrupt [33]. We illustrate the phenomenon of saturation by simulation of the accumulation of pairwise incompatibilities (Figure 5). These simulations (carried out in R; [21]) simply count the number of pairwise incompatibilities between two lineages as substitutions fix, allowing each new substitution to be incompatible with any of the existing ones at that time (with probability *p*). As Figure 5 shows, incompatibilities (*i.e*., interactions that affect hybrid fitness) in saturated systems continue to snowball while the number of individual loci contributing to hybrid incompatibility starts to increase linearly, breaking the relationship between sterility-affecting loci and DMIs. Our simulations are consistent with recent analytical results that also show that, as *p* increases, the number of incompatibility loci tends to underestimate the number of pairwise incompatibilities [33].

**Figure 5:**
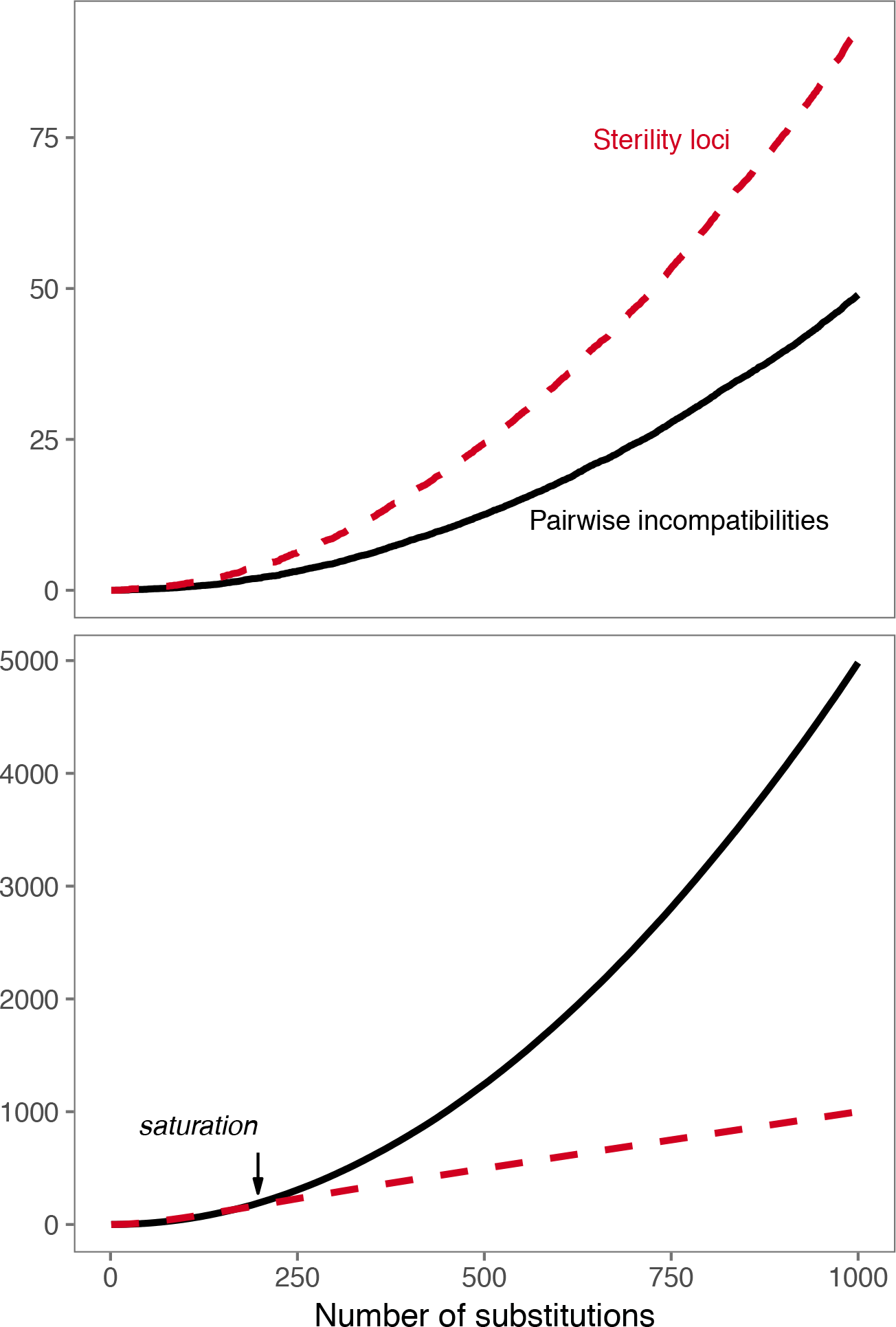
The number of pairwise incompatibilities (showing the “snowball” effect, solid lines), and the number of loci involved in hybrid incompatibilities (red dashed lines), for two cases that differ in the per-substitution probability of pairwise incompatibility (above, *p* = 0.0001; below, *p* = 0.01). Above: In systems that have not saturated, the number of incompatibilities and the number sterility loci grow at a similar rate. While the number of sterility loci is roughly twice the number of incompatibilities, both quantities grow at a faster-than-linear rate. Below: After saturation (around 200 substitutions), the number of loci involved in incompatibilities increases linearly, no longer tracking the number of DMIs. Values are the mean of 100 simulations for each case.

Saturation of the gene network for pollen fitness would be consistent with our observation that several pollen-sterile ILs showed multiple interactions (which suggests they are involved in a higher-order DMI or multiple pairwise ones) and with the previously observed lack of snowball for this phenotype ([18]. Interpreted under a framework of saturation, these two observations suggest many loci underlying pollen fertility are already involved in at least one incompatible interaction between species, so that the accumulation of new interactions does not result in the increase of unique sterility loci at a similar rate. In other words, pollen sterility might be close to saturation for hybrid incompatibility loci (the nodes in our network; Figure 4), even while the DMIs in which they are involved (the edges in our network; Figure 4) continue to snowball among incompatibility loci. If pollen fertility is indeed saturated, it follows that pollen-affecting DMIs may be accumulating in tomatoes faster than previously thought (*i.e*., [18]) because the number of incompatibility QTL underestimates the number of DMIs. This inference is similar to that of recent simulation work on the accumulation of DMIs in an RNA-folding model, which showed that the number of pairwise DMIs will not snowball in systems where these interactions are converted into higher-order DMIs (when new substitutions result in the involvement of a new locus in an existing DMI; [33]). Interestingly, this recent theoretical work also showed that the rate of accumulation of pairwise DMIs slows down as the probability of incompatibility increases [33]. This implies that the high connectivity of the pollen network could be responsible for the observed lack of snowball even if the network is not fully saturated.

Properties of the genome-wide gene network, such as modularity and degree distribution, could also contribute to our proposed saturation of pollen sterility. For instance, highly modular genomes can allow for high *p* within a module (*e.g*., among the genes controlling pollen fertility) even when there is a low genome-wide mean probability of incompatibility. Previous estimates of genome-wide *p* in tomatoes (∼10^−9^;[18]) are smaller than those estimated for *Drosophila* (10^−7^ − 10^−8^, [3]), and much smaller than would seem to be required for saturation. However, our inference of saturation in the pollen network implies large *p* among pollen loci only, which need not be reflected in *p* genome-wide. A skewed degree distribution could also contribute to an apparent saturation by having highly connected loci involved in multiple DMIs while most loci are not involved in an incompatibility. In other words, pollen sterility could be primarily controlled by highly connected genes (*i.e*., be network ‘hubs’), which have a much higher probability of interacting than the genome-wide average. On the other hand, evolutionary processes could have accelerated the rate of divergence specifically at loci involved in fertility traits (that is, have elevated *K* at pollen loci only, rather than across the whole genome). For reproductive traits, this coevolution could arise from antagonistic male-female interactions during reproduction (*e.g*., [34, 35]).

In contrast to these inferences from our pollen data, our observations imply that seed sterility loci in these species might be more likely to engage in DMIs that involve simple pairwise interactions (rather than higher-order epistasis), and that this phenotype is further from saturation. That is, each additional seed sterility-causing substitution is more likely to originate a new DMI with a unique interacting partner, rather than engaging in fitness-affecting interactions with existing sterility loci. In this case, seed sterility loci themselves will still appear to snowball with divergence between lineages; indeed, seed sterility loci have been shown to snowball between *Solanum* species [18].

While our conjecture regarding the saturation of different sterility phenotypes needs to be confirmed with further work, a more general implication of our results is that QTL mapping (or any count of individual hybrid incompatibility loci) might not always provide a suitable estimate of the number and complexity of accumulating DMIs. Despite the considerable additional experimental burden this implies, future empirical tests of the snowball effect should aim to assess both the number of apparent incompatibility loci and evidence for non-independence among these loci. Moreover, we note that the high frequency of interactions with antagonistic effects on sterility also has implications for the pattern of accumulation of reproductive isolation phenotypes (rather than loci or DMIs) between these species. The amount of reproductive isolation is expected to snowball if DMIs are independent and have, on average, similar fitness effect sizes. If, however, antagonistic epistasis among sterility loci is common, the accumulation of reproductive isolation is expected to be less-than-linear, because new DMIs will tend to contribute ‘diminishing returns’ on fitness. This lack of a snowball specifically in incompatibility phenotypes has been previously noted [36, 37], and pervasive phenotypic antagonism could contribute to explaining this common observation [33].

Finally, in contrast to its phenotypic consequences, the mechanistic basis of pervasive genetic antagonism is unclear. This pattern of diminishing deleterious effects is consistent with observations in *E. coli* [38] and yeast [13], but the underlying basis is unresolved. For *Drosophila*, Yamamoto et al. [39] proposed that similar patterns of antagonism might be expected among loci that are involved in stabilizing selection within populations; however, this is unlikely to be relevant here both because our interacting loci are not normally segregating within populations, and because fertility traits are less likely to be subject to stabilizing selection at some intermediate phenotypic optimum (compared to other phenotypic traits). Alternatively, less-than-additive fitness effects can result when different mutations each have deleterious individual effects within the same developmental pathway, but their pairwise combination suppresses or ameliorates these deleterious effects (as has been observed, for example, with double deletion strains in yeast; [13]). Our proposal that, in saturated systems, new substitutions are likely to interact with loci already involved in hybrid incompatibility, is consistent with sterility-affecting mutations accumulating in a tightly-connected network such as a specific developmental pathway. Of course, additional empirical work is needed to evaluate this possibility. Moreover, the frequency of antagonism and the potential for phenotypic diminishing returns are likely to be dependent on the properties of the diverging gene network. Further theoretical work is needed to determine how specific network properties, such as its connectivity and modularity, can affect the patterns of accumulation of incompatibilities and reproductive isolation.

## Acknowledgements

We thank Ata Kalirad and Ricardo Azevedo for helpful discussion, and our three reviewers for valuable comments. This work was supported by National Science Foundation awards DEB-0532097 and DEB-0841957 to LCM. C.D. Muir was supported by an NSF Graduate Research Fellowship. Research assistance was provided by T. Nakazato, E. B. Josephs, M. Yakub, E. Lines, and K. Wolt.

---

**Figure S1:**
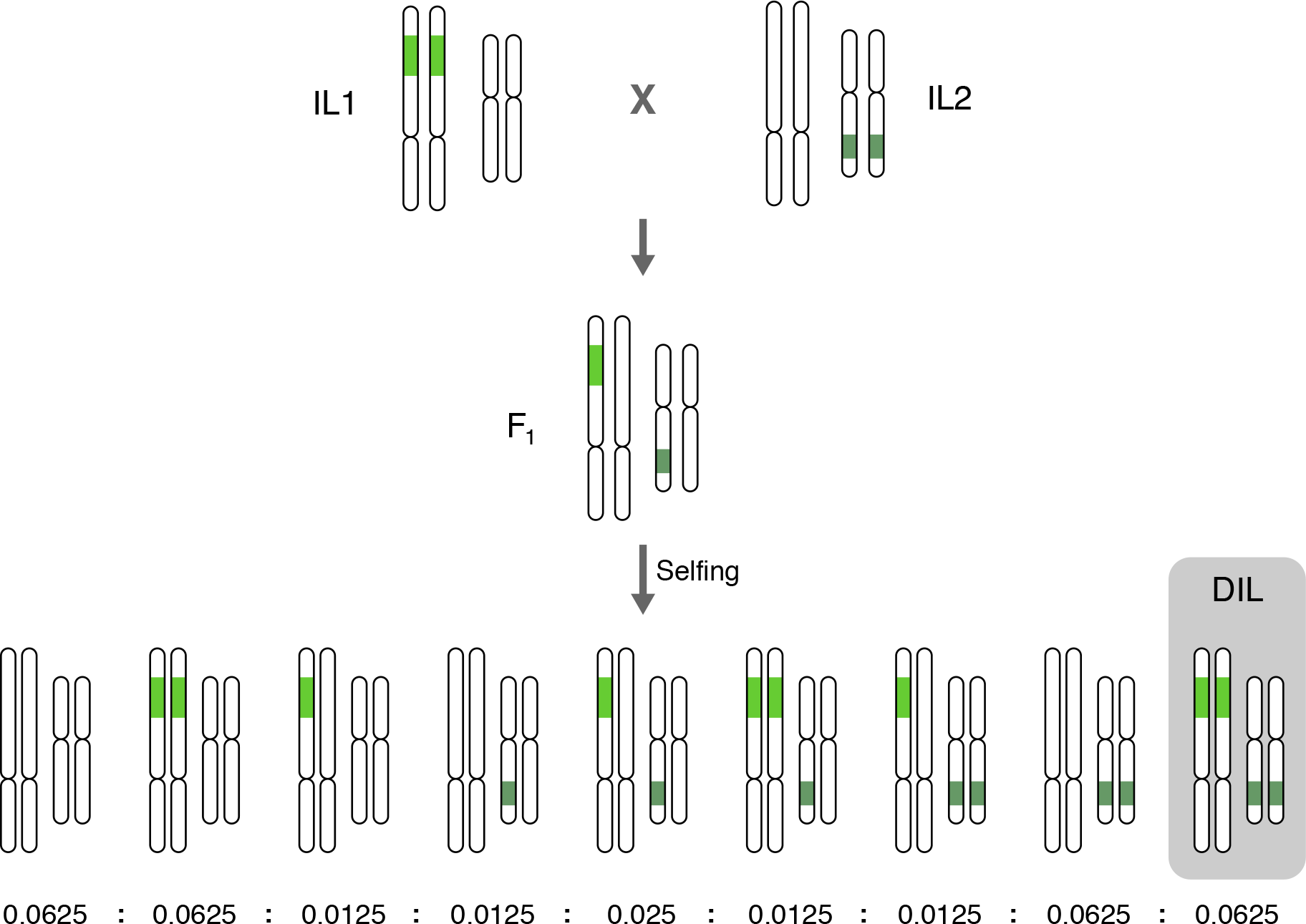
Diagram of cross made to generate the double-introgression lines (DILs). Green shaded areas within a chromosome represent introgressed *hab* regions in an isogenic *lyc* background. Included are the expected genotypic ratios in an F2 population for each DIL family, if marker transmission is Mendelian.

**Figure S2.**
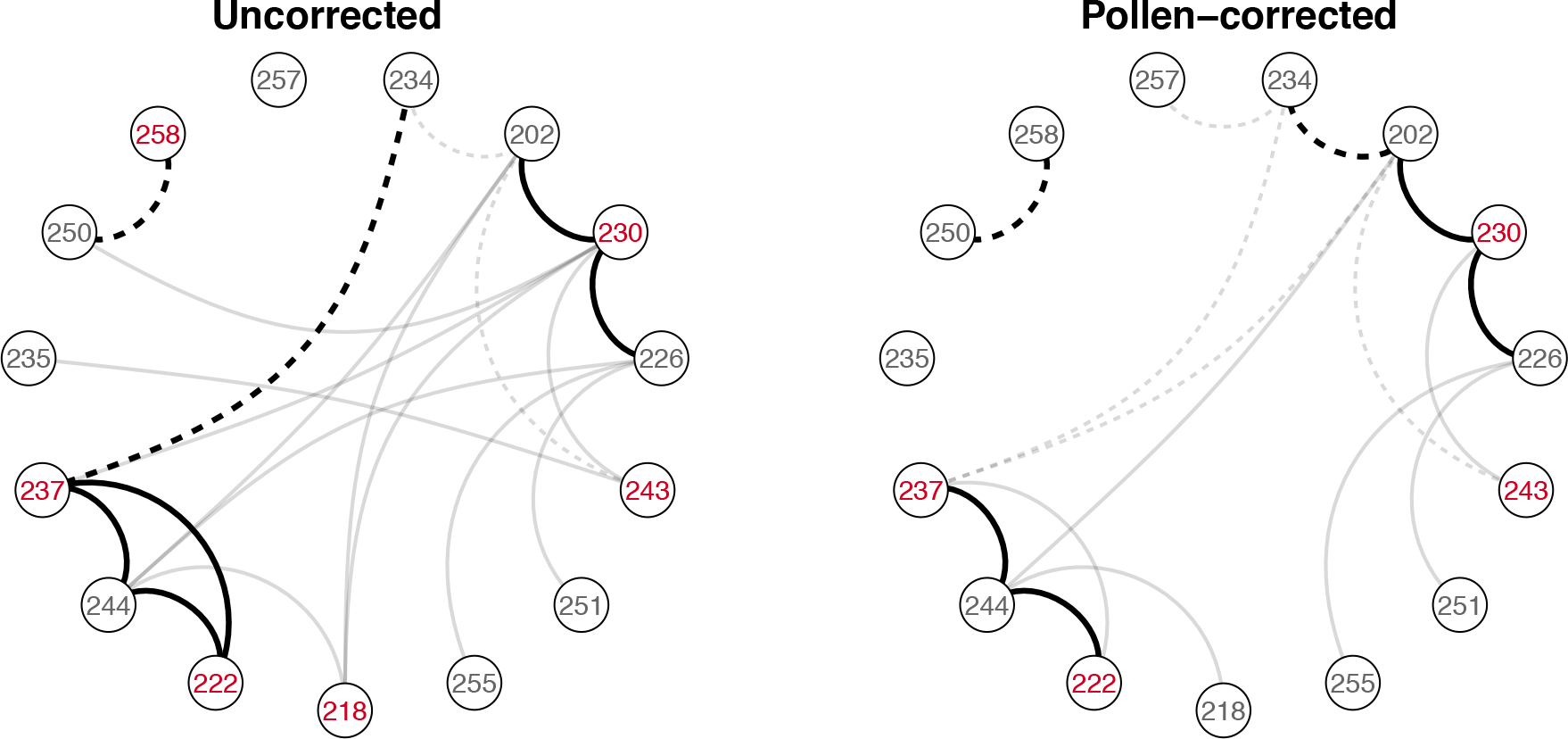
Network representation of interactions in uncorrected self-seed set (left) and pollen-corrected seed (right). Parental ILs (nodes; red label= sterile IL) are connected if they showed a highly significant interaction (FDR=1%). Solid lines denote antagonistic interactions; dashed lines mean synergistic interactions.

**Figure S3.**
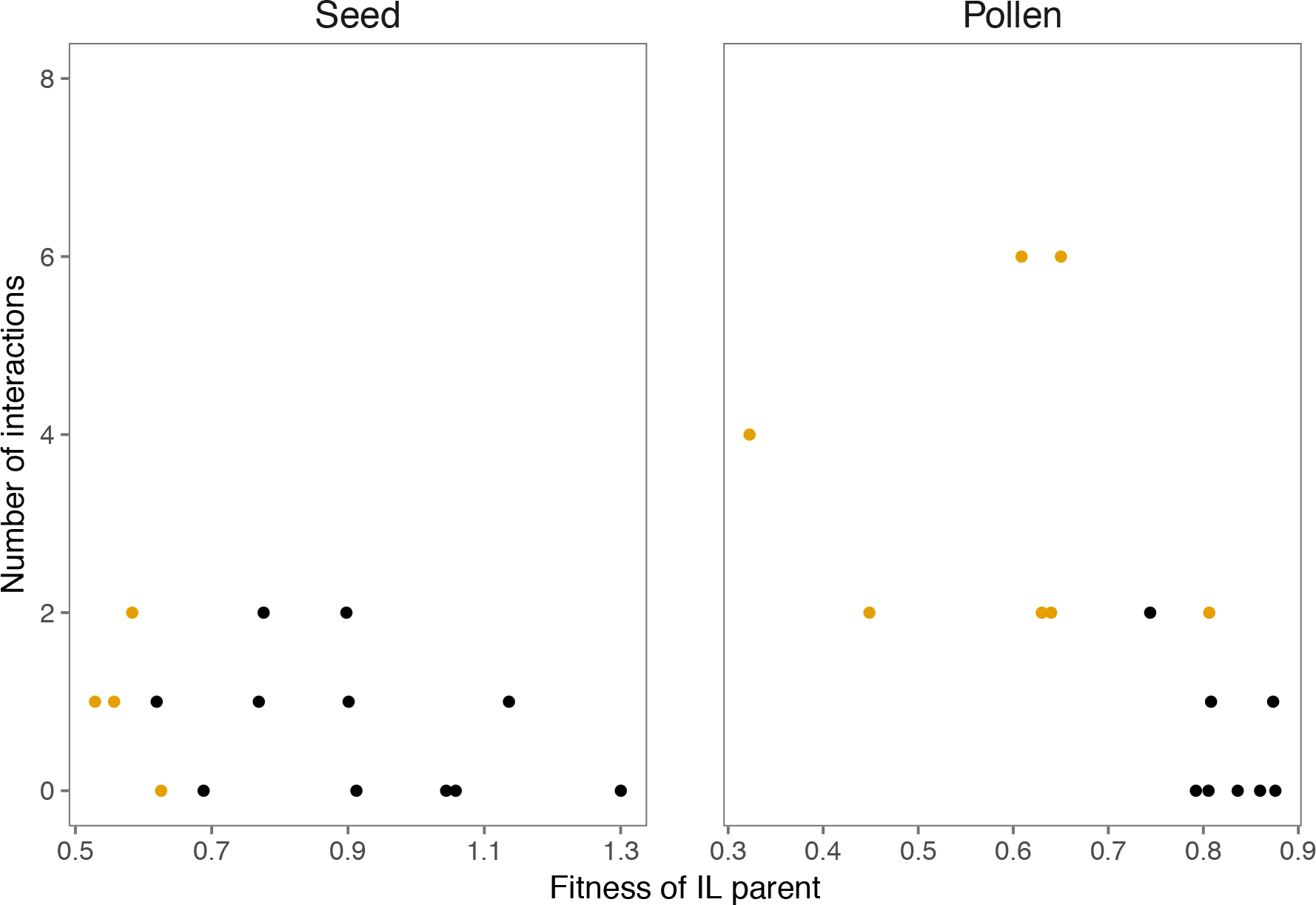
The relationship between parental IL fitness on the number of interactions shown in seed and pollen sterility phenotypes. In pollen, ILs with larger effects on fitness tend to show more interactions (Negative-binomial GLM *P*=0.003, pseudo-R^2^=0.13). This does not seem to be the case for seed phenotypes. Yellow circles represent sterile ILs, black circles are ILs with no individual effects on fitness (see Table 1).

**Figure S4.**
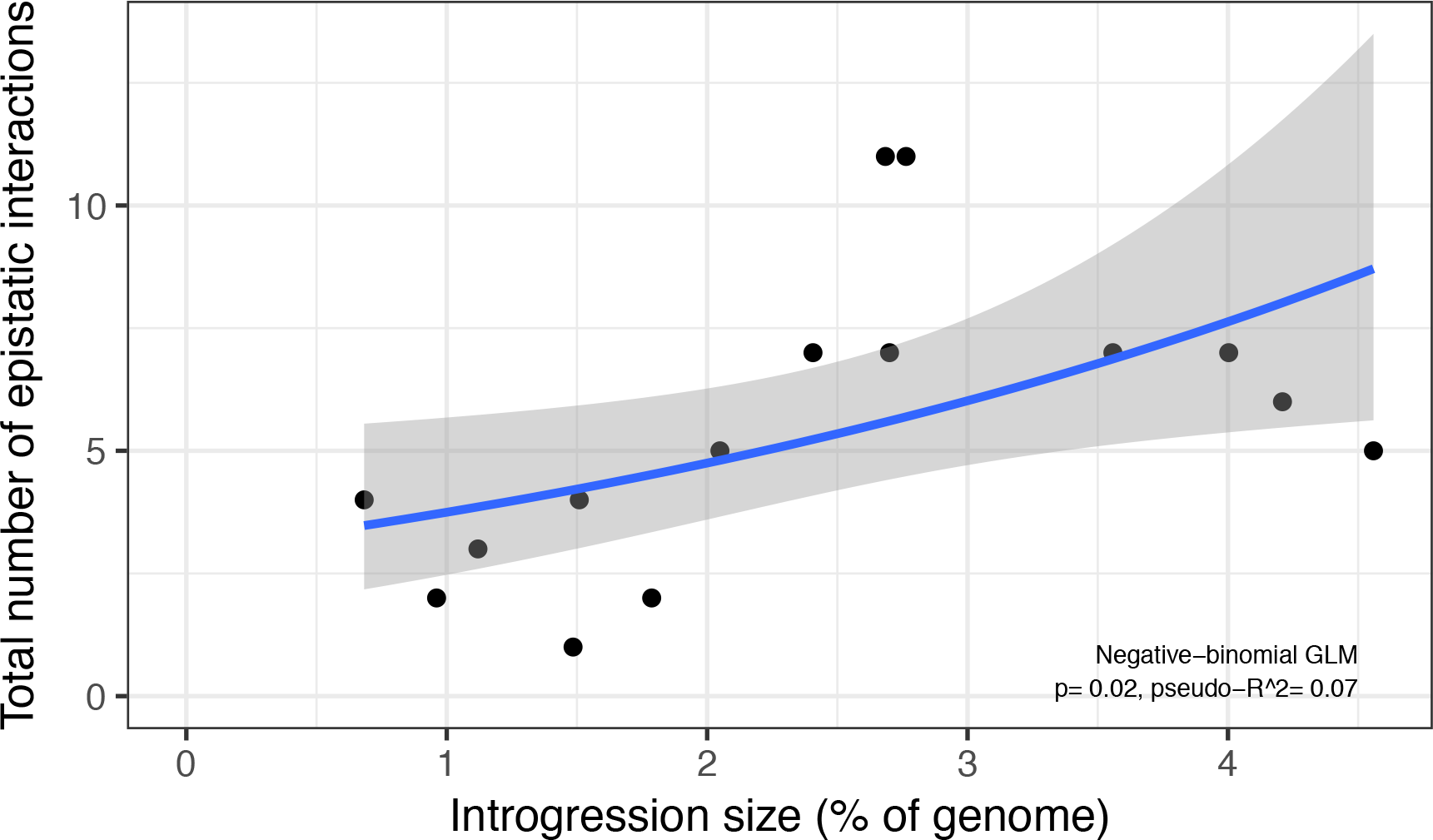
Relationship between the total number of interactions observed for an introgression and its estimated size (as a percent of the *lyc* genome). The total number of interactions is the sum of highly significant (FDR 1%) interactions observed in pollen and seed fertility.

